# Preclinical characterization of immune responses induced by a candidate gonococcal native Outer Membrane Vesicle vaccine

**DOI:** 10.64898/2026.03.09.710637

**Authors:** Ilaria Onofrio, Sthefany Pagliari, Ayah Francis, Molly E. Quinn, Thomas Belcher, Senuri Dissanyake, Clement Twumasi, Iason Vichos, Lukasz A. Grudzein, Christine S. Rollier, Calman A. MacLennan

## Abstract

**Background:** *Neisseria gonorrhoeae* poses significant public health challenges due to multidrug-resistant gonorrhoeic and severe reproductive health complications of untreated infection. No vaccine is licensed to prevent gonorrhea. However, the meningococcal outer membrane vesicle (OMV)-containing vaccine, 4CMenB, provides moderate cross-protection against gonorrhea. We have recently demonstrated that immunization with gonococcal OMV accelerates clearance of gonococcal infection in mice compared with 4CMenB.

**Methods:** To gain insight into possible mechanisms of protection of gonococcal OMV, we evaluated the immunogenicity of GonoVac, a candidate native OMV (nOMV) vaccine against gonorrhea, in mice and rabbits. Three doses of GonoVac were administered intramuscularly from 0.15 to 5 µg in mice, and four doses were used to immunize rabbits at 50 µg per dose, formulated with or without aluminum hydroxide (Al(OH)_3_). Systemic and mucosal antibody responses were evaluated by enzyme-linked immunosorbent assay (ELISA) and serum bactericidal assay (SBA). Cellular responses were assessed by enzyme-linked immunosorbent spot (ELISpot).

**Results:** Immunization with GonoVac formulated with and without Al(OH)_3_ induced significantly higher levels of gonococcal serum and vaginal IgG, and serum bactericidal antibodies, compared with 4CMenB, which induced no serum killing activity. Serum bactericidal activity of GonoVac correlated with anti-gonococcal IgG and IgG2a levels. Serum IgA levels were minimal. Cellular immune responses were higher in mice receiving GonoVac/Al(OH)_3_ compared with GonoVac alone. Immunogenicity was similar for GonoVac produced in a bioreactor and shake flasks.

**Conclusion:** GonoVac elicits robust and functional immune responses in mice and rabbits compared with 4CMenB, supporting its further development as a promising candidate vaccine against gonorrhea.

**Importance:** Gonorrhea, caused by *Neisseria gonorrhoeae*, remains a significant global health concern, disproportionately affecting populations in low- and middle-income countries (LMICs), particularly women. The emergence of multidrug-resistant strains of *N. gonorrhoeae* has raised the concern of untreatable gonorrhea, underscoring the urgent need for effective preventive measures. Although there is no licensed vaccine against gonorrhea, the meningococcal OMV-based vaccine, 4CMenB, is partially effective against the disease and has been recommended for use in high-risk groups.

In this article, we build on previous findings of enhanced efficacy of gonococcal OMV vaccine candidates compared with 4CMenB in the mouse gonococcal infection model to demonstrate the superior anti-gonococcal immunogenicity of a gonococcal OMV-based candidate vaccine (GonoVac) compared with 4CMenB. GonoVac elicits robust immunity in mice, inducing antibodies that are able to kill gonococci, whereas 4CMenB does not. The findings highlight the potential of GonoVac as a promising vaccine candidate for the prevention of gonorrhea worldwide.

## Introduction

Gonorrhea represents a significant global health burden, with an estimated 82 million cases each year (1). Prevalence rates are notably higher among sexually active young adults in LMICs, particularly in sub-Saharan Africa, Southeast Asia, and parts of Latin America (2–4). The bacterium primarily infects the urogenital tract but can also affect the rectum, nasopharynx, and eyes. While men typically present with symptoms such as dysuria and urethral discharge within a week of exposure, up to 50% of infected women are asymptomatic. In women, untreated infections can ascend to the upper reproductive tract, where they can cause pelvic inflammatory disease, infertility, ectopic pregnancy, and chronic pelvic pain (5).

The remarkable ability of the bacterium to adapt and become resistant to antibiotics has rendered many frontline antibiotics ineffective (6–8). The World Health Organization (WHO) has prioritized the development of a gonococcal vaccine as a critical requirement for curbing disease impact and reducing antimicrobial resistance (AMR) (9). However, vaccine development has been hindered by pathogen antigenic variability and immune evasion mechanisms, lack of protective immunity following natural infection and absence of proven correlates of protection against the disease (10–15).

Recent observational studies have renewed optimism by demonstrating partial cross- protection against gonorrhea conferred by an OMV-based serogroup B meningococcal vaccine, MeNZB, rolled out following an epidemic of meningococcal B disease in New Zealand (16). This cross-reactivity is attributed to genetic and structural similarities between *Neisseria gonorrhoeae* and *Neisseria meningitidis*, which share 80-90% genome sequence identity, particularly in outer membrane components (17). A licensed meningococcal group B vaccine, comprising MeNZB OMV and three recombinant proteins (4CMenB), has been shown to reduce gonorrhea incidence by 35-39% among vaccinated individuals (18–23), supporting OMV as a promising approach for gonococcal vaccine development.

4CMenB was recently recommended by the Joint Committee on Vaccination and Immunization in the UK (JCVI) for the prevention of gonorrhea in high-risk groups (24, 25), and subsequently made available as part of the UK national immunization program, for gay and bisexual men with a recent history of multiple sexual partners or a sexually transmitted infection (STI) (26, 27).

We are developing a gonococcal candidate vaccine based on native OMV (nOMV) from *Neisseria gonorrhoeae* with deleted *lpxL1* gene (28). nOMV are naturally shed from Gram-negative bacteria, enriched with lipooligosaccharide (LOS), outer membrane proteins, and lipoproteins, which confer potent immunostimulatory and intrinsic adjuvant properties (29, 30). OMV vaccines are typically formulated with Al(OH)_3_ to attenuate reactogenicity (31, 32). We have recently demonstrated that gonococcal OMVs formulated with Al(OH)_3_ accelerate clearance of gonococcus in the β-estradiol mouse model of vaginal infection (28).

Here, we evaluate the immunogenicity of GonoVac compared with 4CMenB to understand what immunological mechanisms might underlie the enhanced efficacy of gonococcal OMV compared with 4CMenB. We examine GonoVac produced in a bioreactor or shake-flasks, to compare small scale production with a controlled system designed for optimized large-scale bacteria growth, formulated with and without Al(OH)_3_. Immune parameters assessed include gonococcal-specific systemic and mucosal IgG and IgG subclasses, serum IgG avidity, IgA and IgM, antibody-dependent complement-mediated killing of gonococci, and induction of cytokine-producing lymphocytes.

## Results

To investigate the immunogenicity of GonoVac, groups of six BALB/c mice were immunized intramuscularly with four different doses (0.15, 0.5, 1.5 or 5 μg protein-equivalent) of bioreactor- or 5 µg of shake-flask-produced GonoVac formulated with or without Al(OH)_3_. Five µg of 4CMenB (OMV protein-equivalent) or Al(OH)_3_ alone were administered to two further groups, and one group received no immunizations. Each mouse received 3 immunizations given 3 weeks apart and blood samples were collected 3 weeks after each immunization (at week 3, 6 and 9) (**Figure 1**).

**Figure 1:**
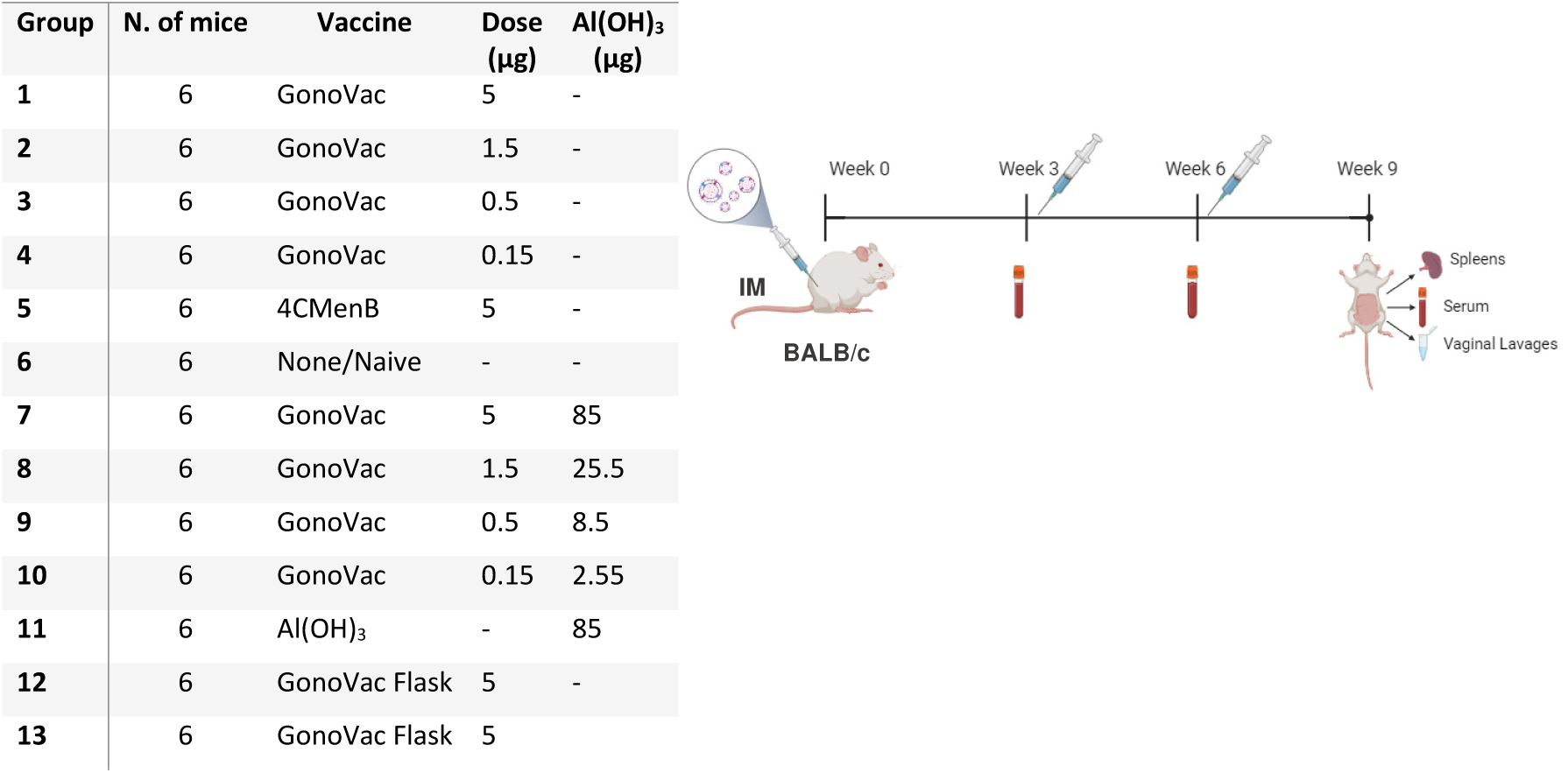
Experimental design to evaluate the immunogenicity in mice of GonoVac in a dose escalation study with and without Al(OH)_3_. Groups of six 6–8-week-old BALB/c mice were immunized intramuscularly with doses of GonoVac ranging from 5 µg to 0.15 µg (total protein content) produced in bioreactor or 5 µg produced in shake flask, formulated with or without Al(OH)_3_. The 4CMenB control group received 5 µg of the dOMV protein equivalent. Naïve mice served as a non-immunized control group, while the Al(OH)_3_ negative control group received adjuvant only. Each group received 3 immunizations of the assigned vaccine at 3-week intervals (week 0, week 3, week 6), and blood samples were collected 3 weeks post each dose at weeks 3, 6 and 9. Spleens and vaginal lavages were collected post-mortem at week 9.

### GonoVac induces higher levels of serum gonococcal IgG compared with 4CMenB

GonoVac (0.5–5 µg), formulated with or without Al(OH)_3_, induced significantly higher levels of serum anti-gonococcal IgG compared with 4CMenB, unvaccinated mice or mice receiving Al(OH)_3_ alone after one, two or three immunizations (p<0.0001 for all comparisons, Two-way ANOVA; **Figure 2A**).

**Figure 2.**
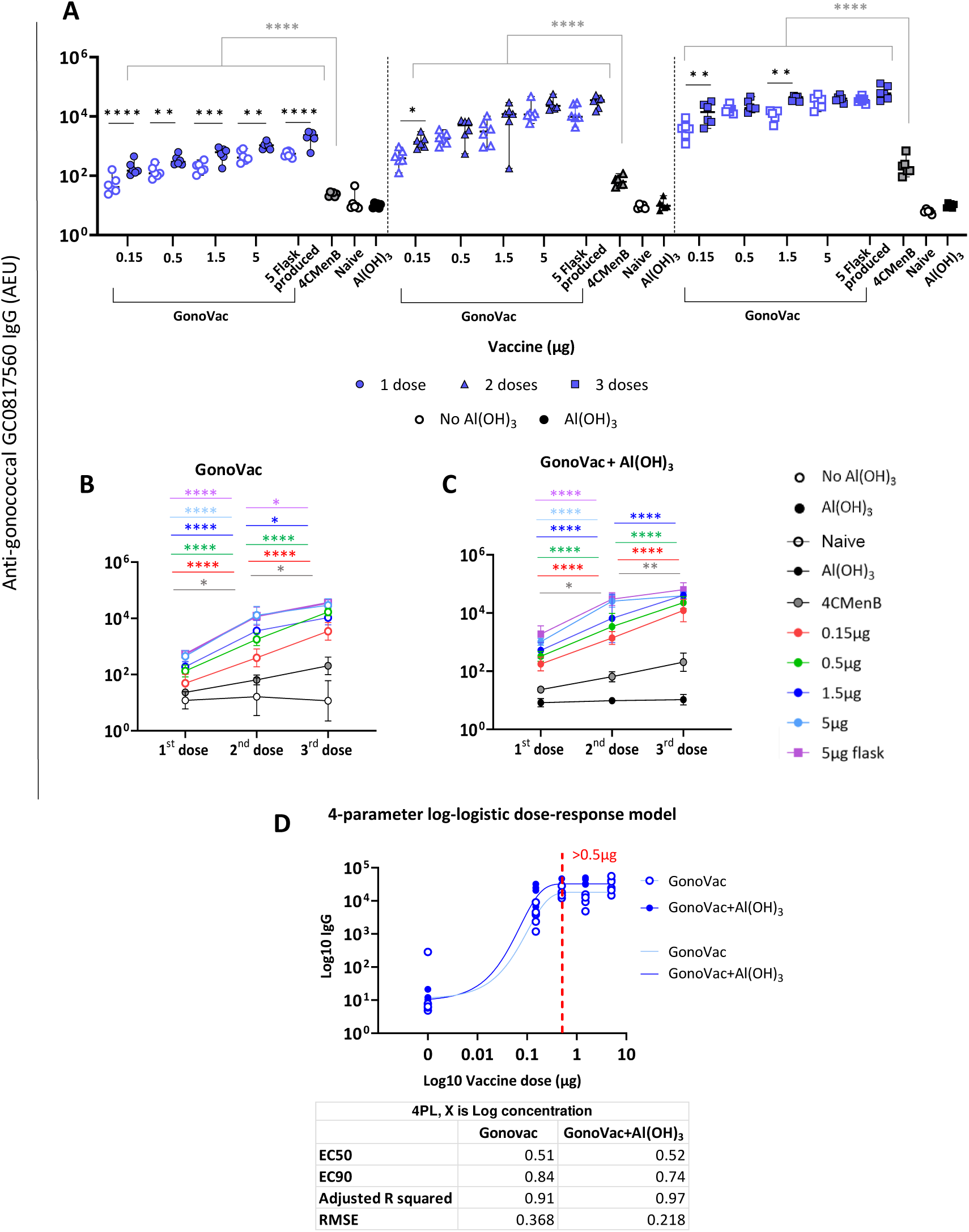
Serum anti-gonococcal IgG responses following a dose escalation of GonoVac with and without aluminum hydroxide (Al(OH)_3_) in mice. 6-8 week-old BALB/c mice were immunized with escalating doses (0.15-5 μg) of GonoVac produced in bioreactor or 5 μg produced in shake flask, with and without Al(OH)_3_ or 4CMenB or Al(OH)_3_ only. Naïve mice were not immunized. Mice received 3 doses at 3-week intervals (days 0, 21, and 42). Serum samples were collected 3 weeks post each dose and analyzed for anti-gonococcal IgG responses against lysates of the GC_0817560 strain using a standardized ELISA. (A) Anti-gonococcal IgG responses - individual mice are represented by the circles (after 1 immunization), triangle (after 2 immunizations) and squares (after 3 immunizations) (median ± 95% CI; n=6). Top bars with stars in grey indicate the significance compared to 4CMenB. (B-C) are the timelines of the vaccine-induced IgG responses (B) GonoVac alone; (C) GonoVac/Al(OH)_3_ (median ± 95% CI; n=6). Statistical test 2way ANOVA with Šídák’s correction for multiple comparisons. P values * p<0.05, ** p<0.01, *** p<0.001, **** p<0.0001. (D) GonoVac-induced anti-gonococcal serum IgG dose-response curve following 3 doses of GonoVac (light blue) or GonoVac/Al(OH)_3_ (blue). Dose-response curve generated by fitting a 4-parameter log-logistic model. Y-axis are Log10 of the anti-gonococcal IgG response; X-axis are Log10 of the vaccine dose administered, with dose 0 replaced by a small value for plotting purposes. The red line indicates where the IgG response plateaus. EC50 – half maximal effective concentration; EC90 – 90% maximal effective concentration; RMSE – root mean square error.

Formulations of GonoVac with Al(OH)_3_ induced higher gonococcal-specific serum IgG responses than equivalent formulations without Al(OH)_3_ after the first dose (p<0.01 to p<0.001). However, there was little difference in IgG levels with and without Al(OH)_3_ following two or three doses of GonoVac (**Figure 2A**). Serum gonococcal-specific IgG levels increased significantly for all GonoVac dose levels and formulations following a second and third immunization, except after dose three at the 5 µg dose level with Al(OH)_3_ (**Figure 2B-C**), suggesting that two 5 µg immunizations are sufficient to achieve a peak anti-gonococcal IgG response.

### Low Doses of GonoVac achieve maximal IgG responses after 3 doses

We evaluated the relationship between GonoVac dose and magnitude of gonococcal-specific serum IgG response following a three-dose immunization regimen, administered with or without Al(OH)_3_. Fitting a 4-parameter log-logistic (PL) regression model to the serum anti-gonococcal IgG levels gave a sigmoidal dose-response relationship. Antibody levels increased with GonoVac doses up to 0.5 µg, beyond which responses plateaued (**Figure 2D**). The adjusted R^2^ exceeded 0.90 for both datasets and the root mean square error (RMSE) values were below 0.5, indicating a high degree of model fit (33) and low model prediction error (34, 35). EC50 (half maximal effective concentration) and EC90 (90% maximal effective concentration) (36) were also calculated, with the EC₉₀ indicating that less than a 1 µg dose is sufficient to elicit a near-maximal antigen-specific serum IgG response (**Figure 2D**).

### GonoVac promotes broad IgG subclass switching

Gonococcal-specific serum IgG1, IgG2a, IgG2b, and IgG3 subclasses analysis by ELISA following three immunizations suggests that GonoVac and 4CMenB primarily induce an IgG1 isotype response in mice (∼4 log_10_ increase in IgG1 from baseline; **Figure 3A**). Levels of serum anti-gonococcal IgG1, IgG2a, IgG2b and IgG3 elicited by GonoVac were significantly higher than those induced by 4CMenB independent of dose (**Figure 3A-D**).

**Figure 3:**
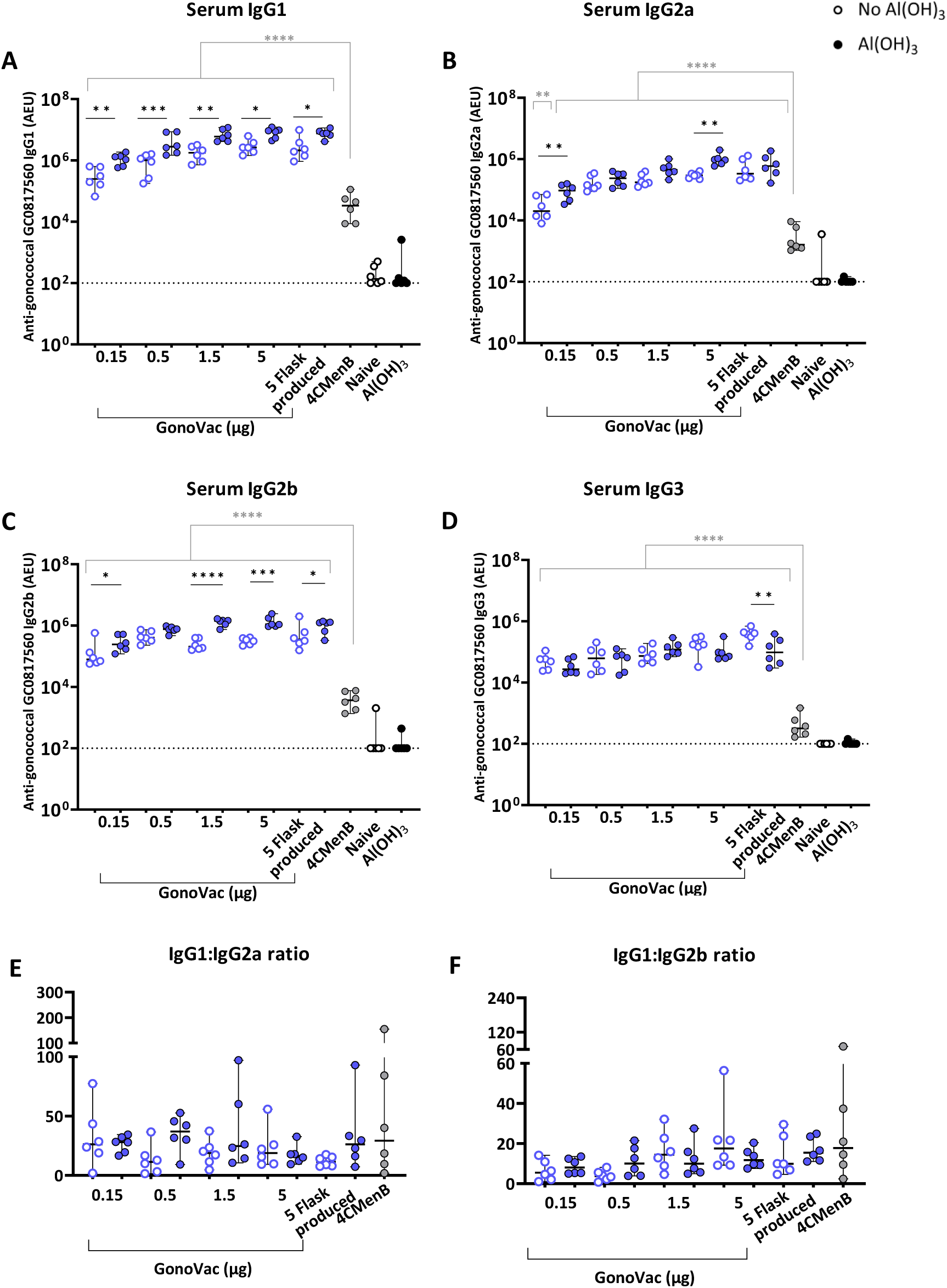
Serum anti-gonococcal (A) IgG1, (B) IgG2a, (C) IgG2b and (D) IgG3 responses following a dose escalation of GonoVac with and without Al(OH)_3_ in mice. 6-8 week-old BALB/c mice were immunized with escalating doses (0.15-5 µg) of GonoVac produced in fermenter or 5 µg produced in shake flask, formulated with and without Al(OH)_3_, 5 µg of 4CMenB or Al(OH)_3_ alone as control. Naïve mice were not immunized. Mice received 3 doses at 3-week intervals (days 0, 21, and 42). Anti-gonococcal IgG subclasses (A) IgG1, (B) IgG2a, (C) IgG2b and (D) IgG3 were determined in serum samples collected after 3 immunizations by ELISA using lysates of the GC_0817560 strain as antigens. All the results are expressed in Arbitrary ELISA Units (AEU). The horizontal dashed line represents the limit of quantification (LOQ), determined as the AEU of the lowest point of the standard curve adjusted by the minimum dilution factor applied to the samples. Serum IgG1 and IgG2a titers were used to calculate (E) IgG1:IgG2a and (F) IgG1:IgG2b providing insight into the type of immune response generated: a higher ratio suggests a predominant Th2-type response, while a lower ratio indicates a stronger Th1-type response. Individual mice are represented by the circles (median ± 95% CI; n=6). Statistical test 2way ANOVA with Šídák’s correction for multiple comparisons; grey stars indicate the significance compared to 4CMenB. P values: * p<0.05, ** p<0.01, *** p<0.001, **** p<0.0001.

At all dose levels tested, GonoVac formulated with Al(OH)_3_ induced significantly higher levels of serum anti-gonococcal IgG1 compared with GonoVac without Al(OH)_3_ (p=0.001 to 0.5-Two-way ANOVA; **Figure 3A**), consistent with aluminum adjuvants promoting Th2 over Th1 immunity (37). Serum gonococcal-specific IgG2a levels were significantly higher with Al(OH)_3_ formulation at only two dose levels (5 µg and 0.15 µg) (**Figure 3B**), while IgG2b titers were significantly higher with Al(OH)_3_ formulation at three dose levels (**Figure 3C**).

In contrast, anti-gonococcal IgG3 responses were similar between formulations with and without Al(OH)_3_ at all doses of GonoVac, except for 5 µg flask-produced GonoVac which induced more IgG3 when formulated without Al(OH)_3_ (p = 0.002 - Two-way ANOVA; **Figure 3D**). We determined IgG1:IgG2 ratios to provide insight into the type of immune response generated (38, 39). After three immunizations, no consistent differences were observed between Al(OH)_3_-adjuvanted and -unadjuvanted GonoVac groups (**Figure 3E-F**).

Serum anti-gonococcal IgA levels mostly did not differ significantly between GonoVac, 4CMenB, and adjuvant-only controls (**Figure 4A**). At 1.5 and 5 µg doses, GonoVac without Al(OH)_3_ induced significantly higher IgM levels than GonoVac/Al(OH)_3_ (**Figure 4B**). We assessed anti-LOS serum IgG levels and found that GonoVac alone induced significantly greater anti-LOS IgG responses compared to GonoVac/Al(OH)_3_ at 1.5 and 5 µg dose levels (**Figure 5**).

**Figure 4:**
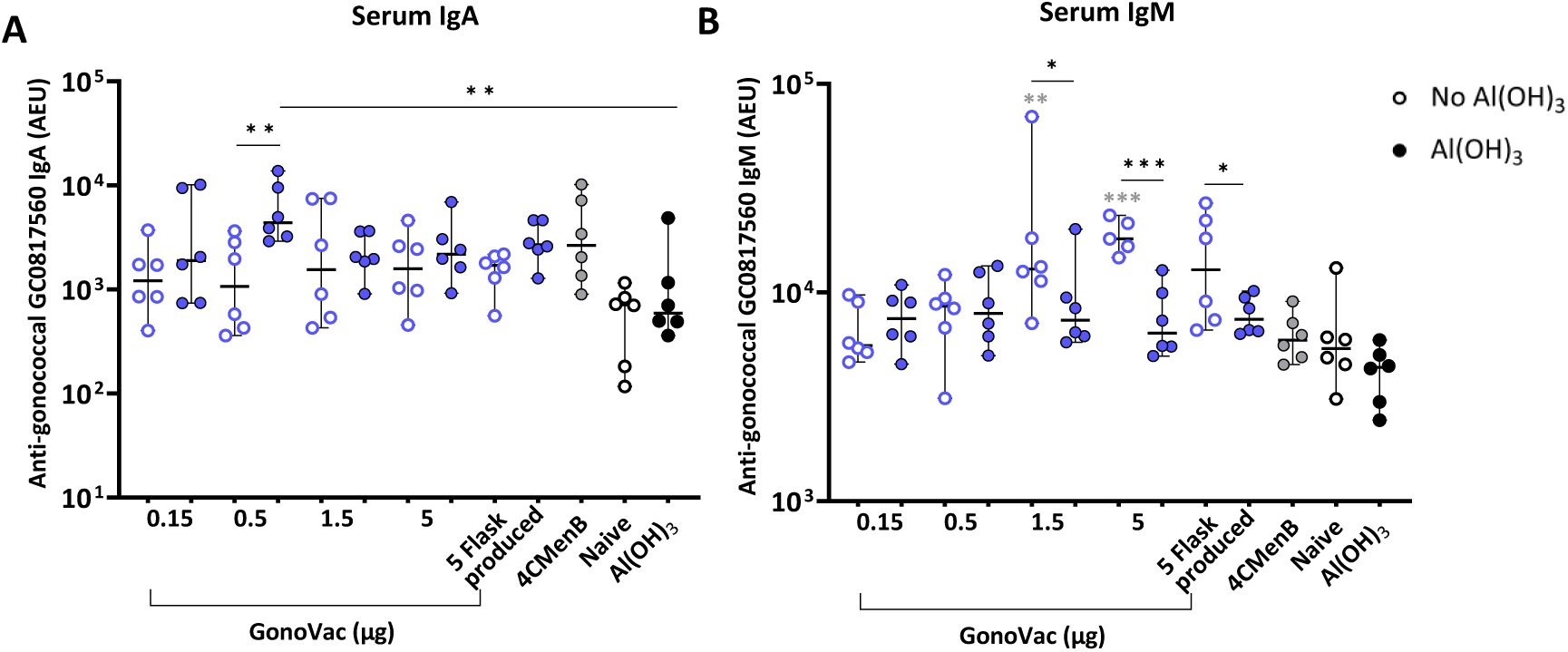
Serum anti-gonococcal IgA and IgM following a dose escalation of GonoVac with and without Al(OH)_3_. 6-8 week-old BALB/c mice were immunized with escalating doses (0.15-5 μg) of GonoVac produced in bioreactor or 5 μg produced in shake flasks, formulated with and without Al(OH)_3_, 5 µg of 4CMenB or Al(OH)_3_ alone as control. Naïve mice were not immunized. Mice received 3 doses 3 weeks apart (days 0, 21, and 42). Anti-gonococcal serum (A) IgA and (B) IgM titers were determined in samples collected after 3 doses. Individual mice are represented by the circles (median ± 95% CI; n=6). All the results are expressed in Arbitrary ELISA Units (AEU). Statistical test 2way ANOVA with Šídák’s correction for multiple comparisons; grey stars indicate the significance compared to 4CMenB. P values: * p<0.05, ** p<0.01, *** p<0.001.

**Figure 5:**
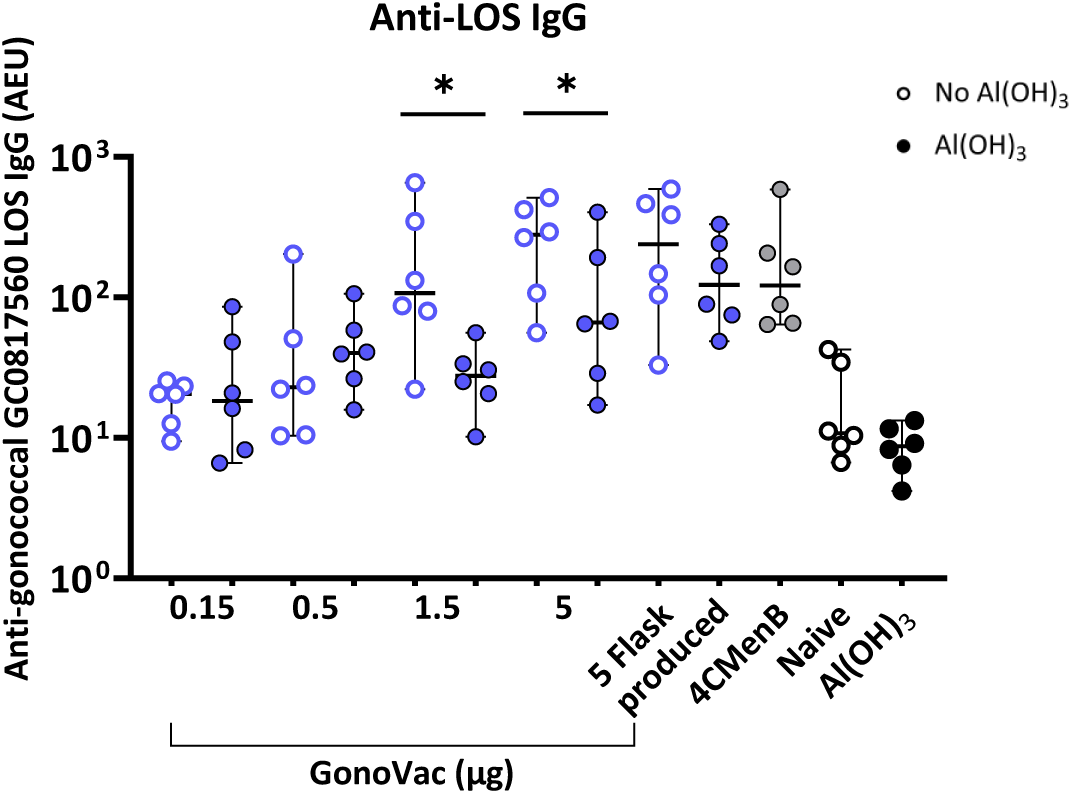
Serum anti-gonococcal LOS IgG induced by a dose escalation of GonoVac with and without Al(OH)_3_. 6-8 week-old BALB/c mice were immunized with escalating doses (0.15-5 μg) of GonoVac produced in fermenter or 5 μg produced in shake flasks, formulated with and without Al(OH)_3_, 5 µg of 4CMenB or Al(OH)_3_ alone as control. Naïve mice were not immunized. Mice received 3 doses 3 weeks apart (days 0, 21, and 42). Anti-gonococcal serum anti-LOS IgG responses were determined in samples collected after 3 doses using an in-house standardized ELISA with gonococcal GC_0817560 hot-phenol extracted LOS as antigen. Individual mice are represented by the circles (median ± 95% CI; n=6). All the results are expressed in Arbitrary ELISA Units (AEU). Statistical test 2way ANOVA with Šídák’s correction for multiple comparisons. P values: * p<0.05.

### immunization with GonoVac induces mucosal IgG responses in the vagina

Significantly higher vaginal IgG titers were observed for all groups immunized with GonoVac compared with 4CMenB (p=0.0001 to 0.05; **Figure 6A**). Just 0.15 µg of GonoVac (unadjuvanted) eliciting significantly higher gonococcal-specific mucosal IgG responses than 5 µg of 4CMenB (p<0.01; **Figure 6A**). In contrast, little mucosal gonococcal IgA was detected in vaginal lavages and the levels did not significantly differ between GonoVac, 4CMenB and negative control groups (**Figure 6B**).

**Figure 6:**
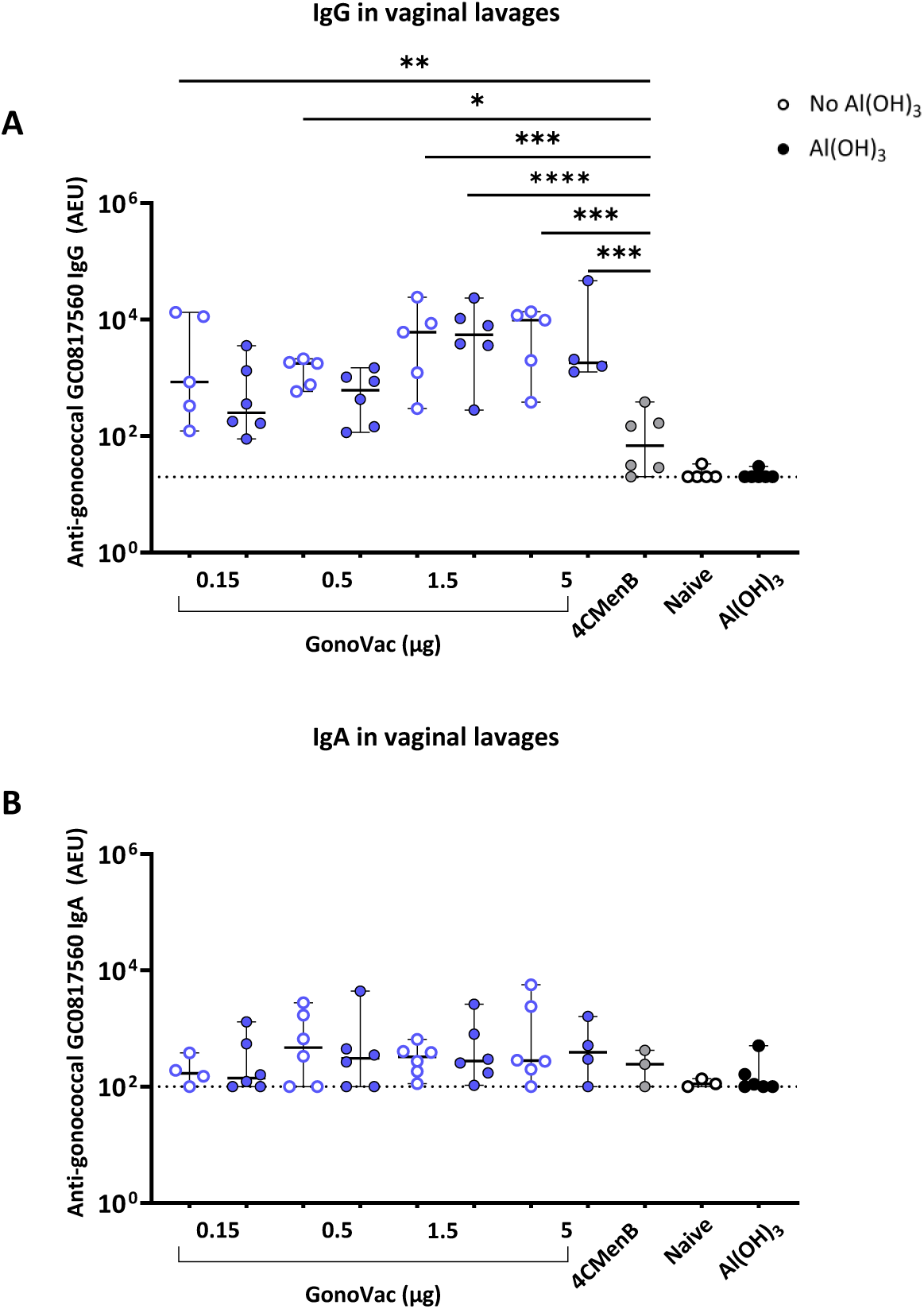
Vaginal anti-gonococcal IgG and IgA responses following a dose escalation of GonoVac with and without Al(OH)_3_. 6-8 week-old BALB/c mice were immunized intramuscularly with escalating doses (0.15-5 µg) of GonoVac, formulated with and without Al(OH)_3_, 5 µg of 4CMenB or Al(OH)_3_ alone as control. Naïve mice were not immunized. Mice received 3 doses at 3-week intervals (days 0, 21, and 42) and vaginal lavages were collected post-mortem at day 63. Mucosal antibody responses were determined by ELISA using lysates of the GC_0817560 strain as antigens: (A) vaginal anti-gonococcal IgG and (B) vaginal anti-gonococcal IgA. Individual mice are represented by the circles (median ± 95% CI; (A) n=5 to 6 – (B) n=3 to 6). Statistical test 2way ANOVA with Šídák’s correction for multiple comparisons. P values: * p<0.05, ** p<0.01, *** p<0.001, **** p<0.0001. All the results are expressed in Arbitrary ELISA Units (AEU). The horizontal dashed line represents the limit of quantification (LOQ), determined as the AEU of the lowest point of the standard curve adjusted by the minimum dilution factor applied to the samples.

### Avidity of gonococcal-specific IgG increases with additional doses of GonoVac

To evaluate anti-gonococcal antibody functionality, we assessed serum IgG avidity and antibody-dependent complement-mediated killing of gonococci. Higher serum IgG avidity was observed with GonoVac alone compared with GonoVac/Al(OH)_3_ following a single 1.5 µg immunization or three 0.5 µg immunizations (**Figure 7A**). There was increased IgG avidity following three doses of 5 µg GonoVac with Al(OH)_3_, and at 0.5 µg GonoVac alone, compared with two doses (**Figure 7B-C**). Despite the lower serum anti-gonococcal IgG levels induced by 4CMenB, the avidity of these antibodies was generally higher than the avidity observed in mice immunized with GonoVac after one and two doses, but not three doses (**Figure 7A**).

**Figure 7:**
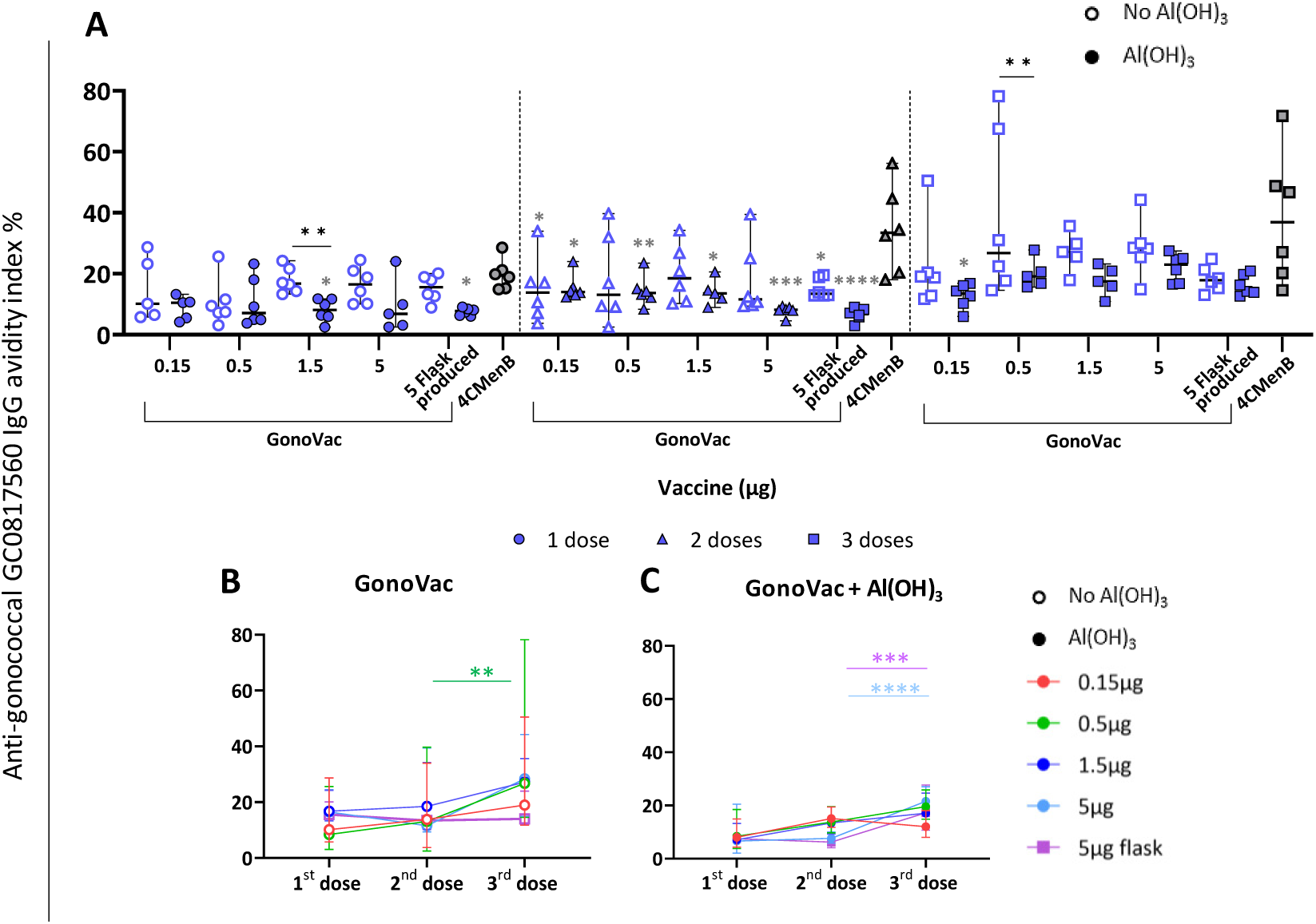
Serum anti-gonococcal IgG avidity and antibody-dependent complement-mediated killing induced by a dose escalation of GonoVac with and without Al(OH)_3_. 6-8 week-old BALB/c mice were immunized intramuscularly with 3 doses 3 weeks apart (days 0, 21, and 42) of escalating doses (0.15-5 μg) of GonoVac produced in fermenter or 5 μg flask-produced, formulated with and without Al(OH)_3_, or 5 µg of 4CMenB. Serum samples were collected 3 weeks post each dose (day 20, 41 and 63). (A-C) Anti-gonococcal IgG avidity was determined in serum samples at each time point using a modified standardized ELISA with lysates of the GC_0817560 strain as antigen and 6 M urea as a chaotropic agent to disrupt low-binding serum IgG antibodies. The avidity was calculated as the percentage of the IgG titer after urea treatment to the titer without urea treatment. Anti-gonococcal IgG avidity responses (A) - individual mice are represented by the circles (after 1 immunization), triangle (after 2 immunizations) and squares (after 3 immunizations). (B) and (C) are the timelines of the gonococcal avidity induced over time by GonoVac (B) without Al(OH)_3_, and (C) with Al(OH)_3_. All graph show median ± 95% CI; n=5 to 6. Statistical test 2way ANOVA with Šídák’s correction for multiple comparisons; grey stars indicate the significance compared to 4CMenB. P values: * p<0.05, ** p<0.01, *** p<0.001, **** p<0.0001.

### GonoVac induces antibody-dependent complement-mediated killing of *N. gonorrhoeae*

The efficacy of antibodies elicited by immunization with GonoVac to kill *N. gonorrhoeae* GC_0817560 was assessed by SBA. GonoVac induced significantly higher SBA titers against GC_0817560 at all dose levels following three immunizations compared with 4CMenB and Al(OH)_3_ alone, as 4CMenB failed to induce bactericidal antibodies (**Figure 8**). Notably, all mice that received GonoVac at doses ≥ 0.5 µg protein-equivalent induced SBA titers ≥4. No significant differences in SBA titers were observed between groups immunized with GonoVac alone compared with GonoVac/Al(OH)_3_.

**Figure 8:**
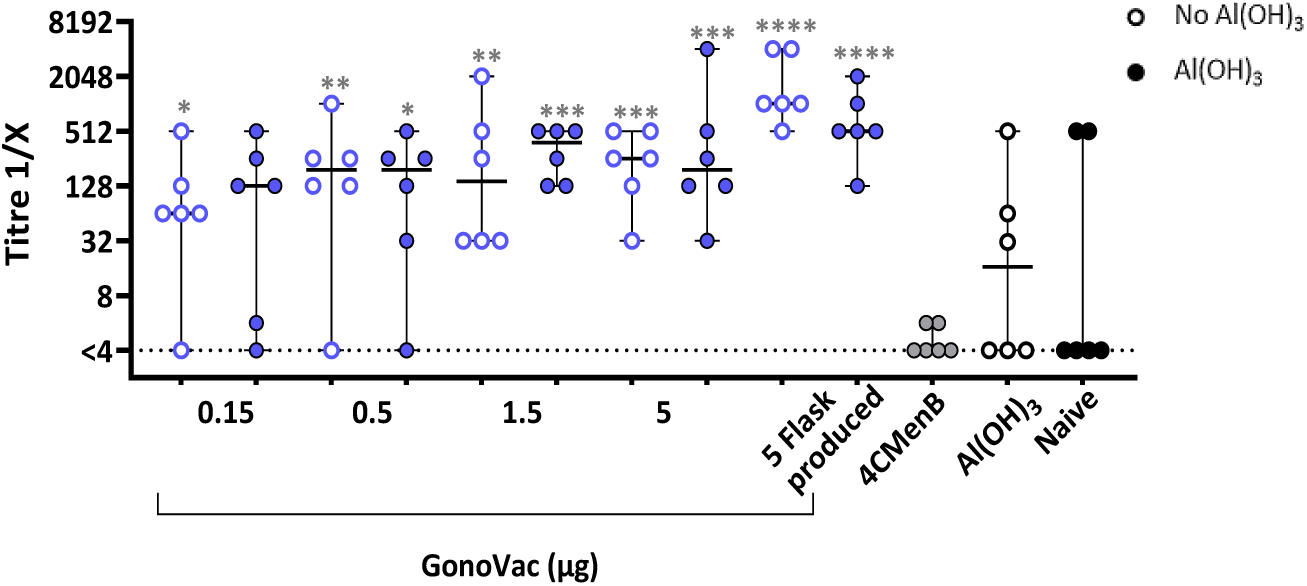
Antibody-dependent complement-mediated killing of *N. gonorrhoeae*. Serum bactericidal assay (SBA) titers against the vaccine parental strain GC_0817560, were assessed in samples collected after 3 immunizations, using an external human source of complement (IgG/IgM/IgA depleted serum). 2-fold serial dilutions of serum were tested, and the titers are represented by the reciprocal of the last dilution of serum inducing >50% killing compared to the complement-alone control. The minimum dilution tested was 1:4. <4 on the graph indicates that no titer was obtained. Individual mice are represented by the circles (median ± 95% CI; n= 6). Statistical test 2way ANOVA with Šídák’s correction for multiple comparisons; grey stars indicate the significance compared to 4CMenB. P values: * p<0.05, ** p<0.01, *** p<0.001, **** p<0.0001.

### Formulation of GonoVac with Al(OH)_3_ induces antigen-specific IFNγ- and IL-17-secreting cell responses in spleens

Cellular responses were evaluated by ELISpot assessing antigen-specific splenocytes secreting IFNγ, IL-4 and IL-17A. Splenocytes from mice immunized with GonoVac and Al(OH)_3_ induced significantly higher numbers of IFN-γ- and IL-17A-secreting cells compared with those receiving GonoVac alone, 4CMenB or Al(OH)_3_ alone (**Figure 9A** and **C**). In contrast, numbers of IL-4-secreting cells were similar between mice immunized with GonoVac with or without Al(OH)_3_ and little increased compared to naïve mice (**Figure 9B**). Although not significant at all dose levels, the median number of IFN-γ- and IL-17A-secreting cells induced by three immunizations with GonoVac without Al(OH)_3_ was higher than that observed in naive mice (**Figure 9A** and **C**). 4CMenB induced-cytokine-secreting cells were not different from those of naïve mice or mice receiving only Al(OH)_3_.

**Figure 9:**
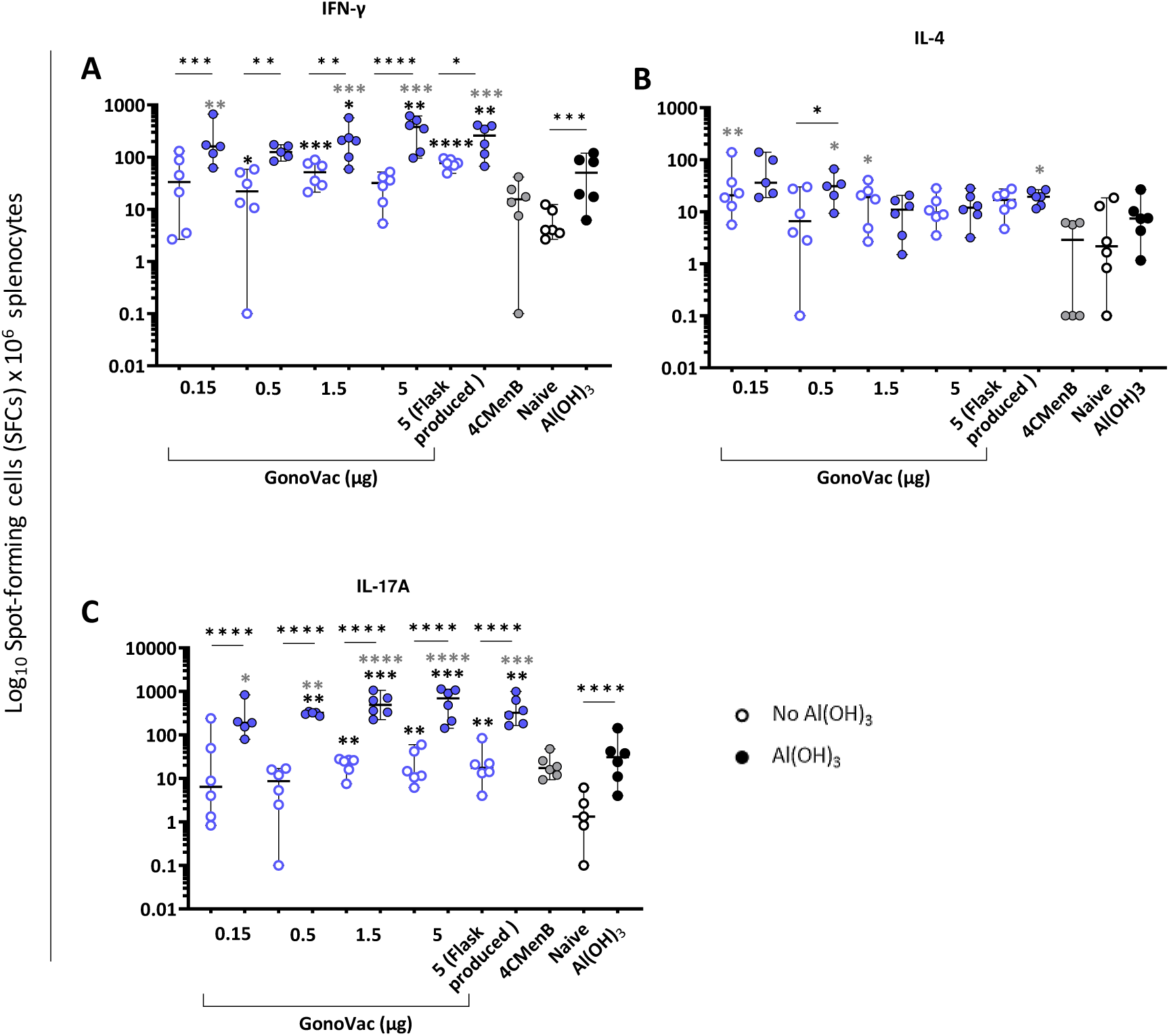
Gonococcal-specific cytokines producing splenocytes induced by a dose escalation of GonoVac with and without Al(OH)_3_. 6–8-week-old BALB/c mice were immunized with a concentration of GonoVac ranging between 0.15 and 5 μg, produced in fermenter or 5 μg-flask derived, with and without Al(OH)_3_. The positive control group received 5 μg of 4CMenB while the negative control groups received Al(OH)_3_ alone or were not immunized (naïve). Mice received 3 doses at 3 weeks intervals (day 0, 21 and 42) and spleens were collected 9 weeks post 1^st^ dose (day 63). Cryopreserved single-cell splenocytes (2.5 x 10^5^ cells) were used to assess vaccine-induced cytokine-producing cells upon restimulation in vitro with 75 μg/mL of gonococcal lysates (GC_0817560) by FluoroSpot assay detecting (A) IFN-γ, (B) IL-4, and (C) IL-17A. Negative control values (unstimulated cells) were subtracted from the corresponding stimulated sample values. Plotted is the log-transformed number of spots forming cells (SFCs) per million of splenocytes. Individual mice are represented by the dots (median ± 95% CI; n=6). Statistical test 2way ANOVA with Šídák’s correction for multiple comparisons; black stars Indicate significance compared with naïve mice while grey stars indicate the significance compared with 4CMenB. P values: * p<0.05, ** p<0.01, *** p<0.001, **** p<0.0001.

### GonoVac produced in shake-flasks and bioreactors induces comparable immune responses in mice

Flask- and bioreactor-produced GonoVac induced equivalent serum IgG, IgG avidity, and cytokine responses, as assessed by Kruskal–Wallis with Dunn’s post hoc test for all comparisons (**Figure 10**), apart from SBA titers which were modestly higher with flask-compared with bioreactor-derived GonoVac when formulated without Al(OH)_3_ (p = 0.01; Figure 10G).

**Figure 10:**
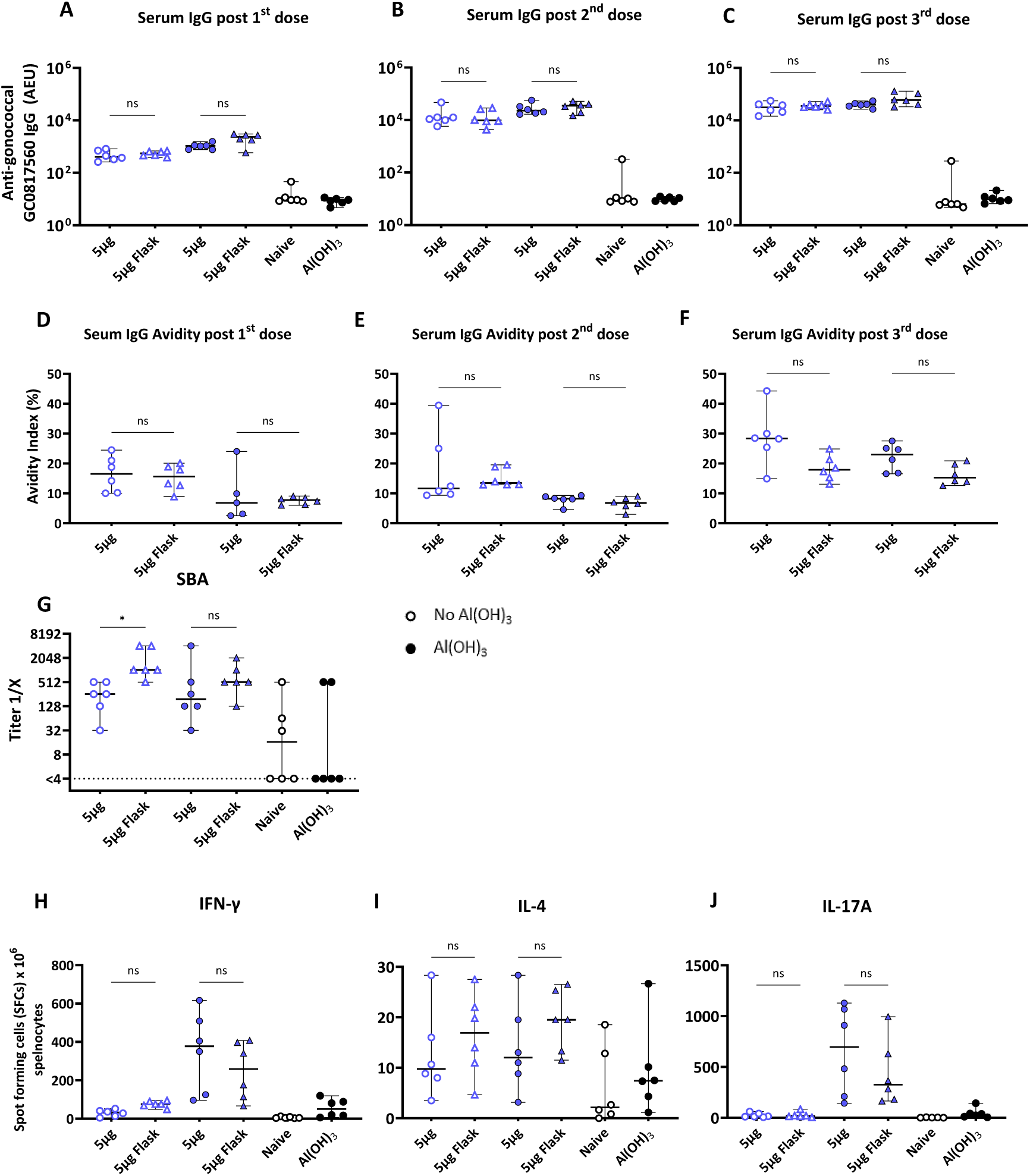
Comparison of the anti-gonococcal serological and cellular responses induced by 5 µg of GonoVac produced in a large or small scale formulated with and without Al(OH)_3_. 6–8-week-old BALB/c mice were immunized with a concentration of GonoVac produced in a 1.5 L fermenter ranging between 0.15 and 5 μg or 5 μg of GonoVac produced in shake flask as control. Negative control groups received Al(OH)_3_ alone or were not immunized (naïve). Mice received 3 doses 3 weeks apart (day 0, 21 and 42) and serum anti-GC_0817560 specific-IgG post (A) 1^st^ dose, (B) 2^nd^ dose (C) post 3^rd^ dose, and IgG avidity post (D) 1^st^ dose, (E) 2^nd^ dose (F) post 3^rd^ dose were assessed by ELISA. Serum and splenocyte samples collected at the end of the study (day 63) were used to assess the functionality of the antibody response in killing the gonococcal vaccine parental strain (GC_0817560) (G) by SBA and (H) IFNγ-, (I) IL-4- and (J) IL-17A-secreting splenocytes. Anti-gonococcal antibody and cellular responses induced by GonoVac produced in small and large scale were compared Dunn’s pairwise multiple comparison with nonparametric Kruskal Wallis test. Individual mice are represented by the dots (median ± 95% CI n=6). P values: * p<0.05; ns - not significant.

### GonoVac-induced SBA titers correlate with serum anti-gonococcal IgG2a

The relationship between serum gonococcal-specific IgG, IgG subclasses, IgG avidity, and SBA titers induced by GonoVac and 4CMenB, was assessed by Spearman’s correlation coefficient (*r*). There is a significant correlation (*r*=0.45, p<0.001) between serum IgG and SBA titers (**Figure 11**). Among the IgG subclasses, IgG2a levels significantly correlate with SBA titers (*r*=0.43, p<0.01), while IgA, IgM IgG1, IgG2b and IgG3 did not correlate. No significant correlations were observed for 4CMenB-induced parameters (n=6; **Supplementary Figure 1**).

**Figure 11:**
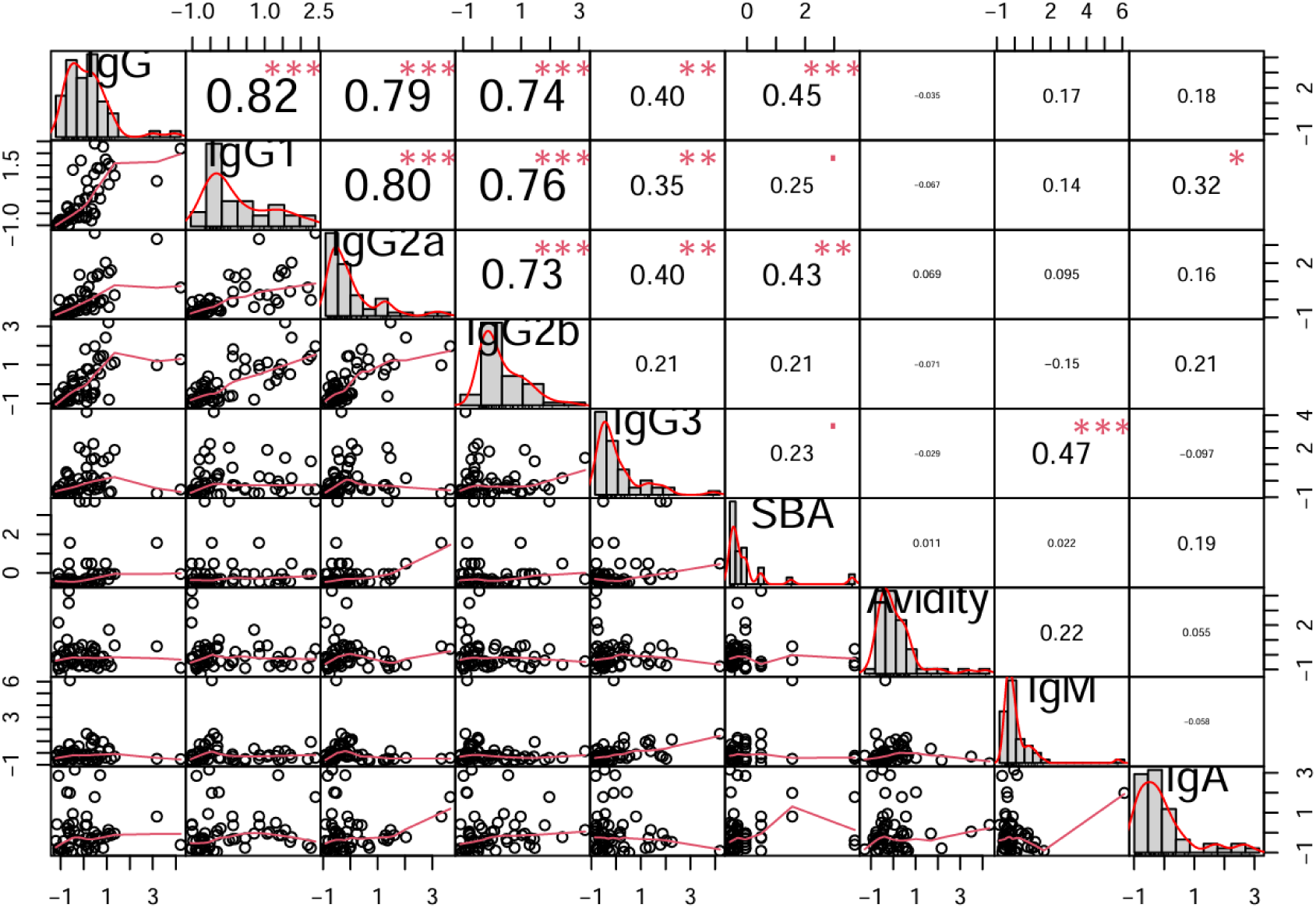
Correlation matrix between GonoVac-induced serum anti-gonococcal IgG, IgM, IgG subclasses, serum bactericidal titers and IgG avidity. Matrix generated utilizing R performance analytics package using data from mice immunized with 3 doses of GonoVac (0.15-5 μg). Diagonally, the histograms represent the distribution of the parameters tested, from top to bottom: serum IgG, IgG1, IgG2a, IgG2b, IgG3, SBA, IgG avidity, IgM and IgA titers. In the upper triangle are shown the correlation coefficients (the bigger the number, the higher the coefficient) with P values (red stars show significance, *p<0.05; **p<0.01; ***p<0.001; red small squares represent correlations that did not achieve significance). In the lower triangle are shown the bivariate scatter plots with a trend line to illustrate relationships between variables.

### Immunogenicity of GonoVac in Rabbits

To evaluate the immunogenicity of GonoVac in rabbits, groups of 6 white New Zealand rabbits (three male and three female per group) were immunized intramuscularly with 4 doses of 50 µg GonoVac with and without Al(OH)_3_ or with Al(OH)_3_ alone. Two groups were included for each formulation. One was culled at day 65 post 1^st^ dose and the other at day 95 (**Figure 12**).

**Figure 12:**
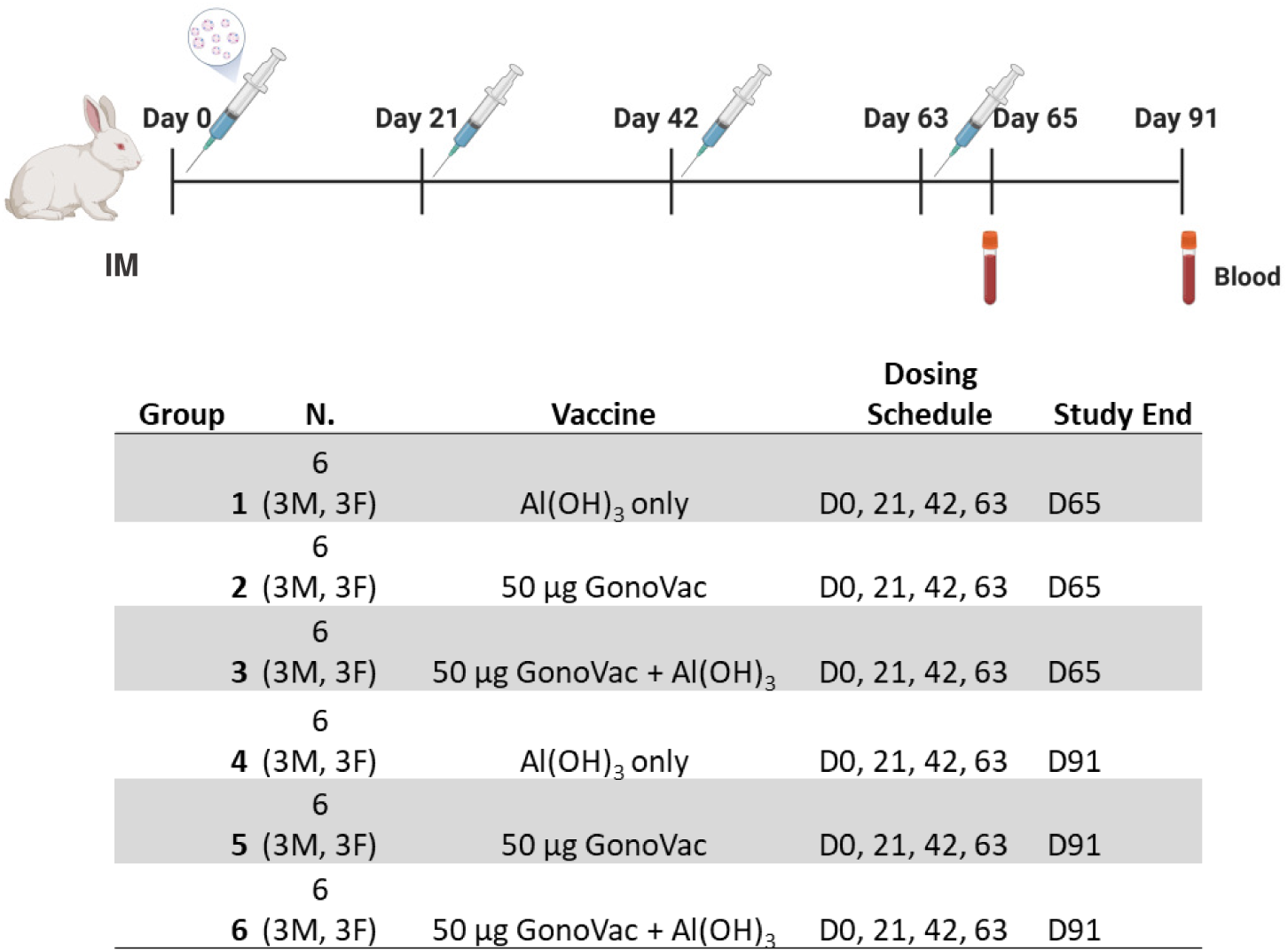
Experimental design to evaluate the immunogenicity in rabbits of GonoVac with and without Al(OH)_3_. Groups of six white New Zealand rabbits (three male and three female per group) received 4 doses of 50 μg GonoVac intramuscularly with and without Al(OH)_3_ or with Al(OH)_3_ alone. Two groups were included for each condition: one group was sacrificed 65 days post 1^st^ dose and the other group 95 days following the 1^st^ immunization.

Serum collected on days 65 and 91 was evaluated for anti-gonococcal IgG antibodies and SBA activity. Both GonoVac and GonoVac with Al(OH)_3_ induced significantly higher serum anti-gonococcal IgG titers compared with placebo controls at both timepoints. There was no significant difference in antibody titers between groups receiving GonoVac with or without Al(OH)_3_ (**Figure 13A**).

**Figure 13:**
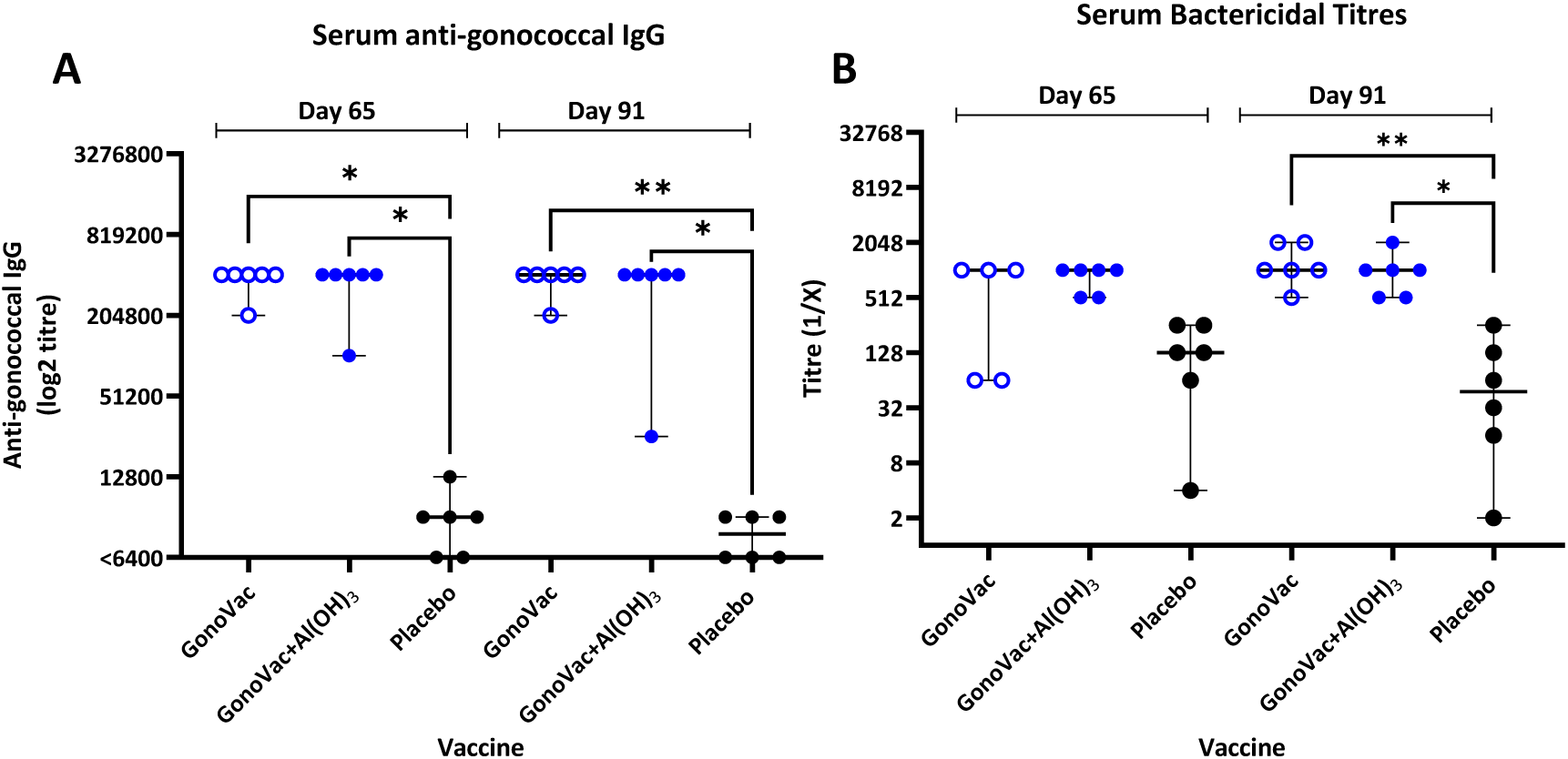
Anti-gonococcal serum bactericidal antibodies in rabbit sera following immunization with GonoVac with or without Al(OH)_3_ or Al(OH)_3_ placebo control. Groups of 6 white New Zealand rabbits (three male and three female per group) were immunized intramuscularly with 4 doses of 50 μg GonoVac with and without Al(OH)_3_ or placebo. Two groups were included for each condition, half were culled at day 65 post 1^st^ dose and the rest at day 95. Serum gonococcal IgG were determined by endpoint titer ELISA using lysates of the gonococcal vaccine parental GC_0817560 strain as antigens. The titer is represented by the reciprocal of the last dilution giving an optical density (OD450) of at least three standard deviations higher than the OD450 of the negative control (a pool of naïve rabbit sera). 6400 is the limit of detection, titers below this value are represented as <6400. Antibody-dependent complement-mediated killing of the gonococcal vaccine parental GC_0817560 strain was assessed by SBA, using an external human source of complement (IgG/IgM depleted human serum). 2-fold serial dilutions of serum were tested and the titers are represented by the reciprocal of the last dilution of serum inducing >50% killing compared to the complement-alone control. Individual rabbits are represented by the dots (median ± 95% CI). Statistical test: Kruskal-Wallis with Dunn’s multiple comparisons post-test: * p<0.05, ** p<0.01.

SBA analysis showed robust antibody-dependent complement-mediated killing in sera from immunized rabbits, with median titers higher than those of placebo controls with statistical significance achieved at day 91. Functional bactericidal activity was similar between GonoVac alone and GonoVac/Al(OH)_3_ groups, suggesting Al(OH)_3_ does not affect anti- gonococcal antibody levels and functionality in rabbits following a 4-dose regimen (**Figure 13B**).

## Discussion

This study provides a comprehensive analysis of the immune responses induced by the candidate gonococcal nOMV vaccine, GonoVac. Beyond confirming robust immunogenicity of the candidate in relation to induction of serum anti-gonococcal IgG compared with 4CMenB, the work explored gonococcal-specific IgG subclass distribution and IgA and IgM levels, serum antibody-dependent complement-mediated killing, serum IgG antibody avidity, mucosal antibody responses, and induction of gonococcal-specific cytokine- secreting cell responses in spleens. These findings provide insights into the breadth and quality of immunity induced by GonoVac in comparison with 4CMenB.

Even low antigen doses (0.15–0.5 µg) of GonoVac are sufficient to induce significant systemic and mucosal IgG responses that are significantly higher than those induced by the licensed meningococcal group B vaccine, 4CMenB. Dose-response modelling confirmed a plateau in serum IgG titers, following three doses of GonoVac, greater than 0.5 µg indicating an antigen-sparing potential for the vaccine. Three immunizations induced the highest immune responses, though with limited incremental benefit beyond two doses at the 5 µg level, indicating potential to meet a two-dose schedules for global gonococcal vaccine implementation, as specified in the WHO Gonococcal Vaccine Preferred Product Characteristics report (40).

GonoVac induced robust functional serum bactericidal responses against *N. gonorrhoeae* GC_0817560, while such responses were absent in mice immunized with 4CMenB. The lack of killing activity following immunization with 4CMenB could related to the strain of *N. gonorrhoeae* used in the assay or the assay itself, since other investigators have found both mouse (41, 42) and human serum (43) to be bactericidal against other strains of gonococcus following immunization with 4CMenB. Although there is no established correlate of protection for gonorrhea, the SBA is an accepted method of assessing functional antibody activity against Gram-negative bacteria and serves as a correlate of protection for invasive meningococcal disease (44).

Consistent with the murine data, rabbits immunized with either formulation of GonoVac exhibited significantly higher anti-gonococcal IgG levels and SBA titers compared with placebo controls, further supporting the functionality as well as cross-species robustness of the immune responses elicited by this vaccine. There was a moderate yet statistically robust association between total serum IgG levels and SBA titers, with IgG2a likely a primary contributor. Importantly, SBA titers were high in the absence of Al(OH)_3_.

GonoVac without Al(OH)_3_ also induced modestly higher serum gonococcal-specific IgM titers compared to the formulations with Al(OH)_3_ and 4CMenB (45). Although GonoVac-induced IgM did not directly correlate with SBA titers, its presence may contribute to the enhanced ability to kill gonococci compared with 4CMenB. Vaccine-induced vaginal IgG was present in the absence of a significant IgA response, typical of parenteral immunization, suggesting antibody transudation from the blood or local IgG production (46). This finding supports the potential utility of a gonococcal OMV vaccine against gonorrhea, given the genital mucosa is the primary site of gonococcal entry. Notably, GonoVac induced stronger mucosal anti-gonococcal IgG responses than the 4CMenB vaccine, further supporting its potential for protection at the site of infection.

Serum IgG antibodies against gonococcal LOS were higher with GonoVac alone compared with GonoVac/Al(OH)_3_ consistent with the concept that formulation with Al(OH)_3_ reduces the availability of LOS to drive an anti-LOS antibody response. The IgG1-dominant subclass profile observed is consistent with a Th2 immune response, particularly when GonoVac is formulated with Al(OH)_3_ (47), though IgG2a and IgG2b induction and IFN-γ/IL-17A-producing splenocytes indicate that Th1/Th17 responses are also induced.

The ability of GonoVac to induce high IgG and SBA levels, not only in mice but also in rabbits, further indicates the immunostimulatory potency of gonococcal nOMVs. These findings are particularly relevant given SBA is a proposed mechanistic correlate of protection of a preclinical gonococcal mimotope vaccine candidate based on the 2C7 LOS epitope (48). The ability of GonoVac to induce robust SBA responses without Al(OH)_3_, supports investigation of the unadjuvanted vaccine in the clinic.

The immunological comparisons undertaken in this study of GonoVac and 4CMenB are key given the recent recommendations in the UK and New York State to make 4CMenB available to individuals at high risk of gonorrhea (27, 49). 4CMenB, which contains meningococcal OMV, offers partial effectiveness in retrospective analyses of clinical trials against gonorrhea and therefore is best considered a stopgap until more effective vaccines are developed (22, 49). Our hypothesis that gonococcal OMV-based vaccines will offer enhanced efficacy against gonococcal infection compared with 4CMenB was recently shown to be correct in the mouse gonococcal infection model using a precursor of GonoVac (28). The finding of enhanced immunogenicity of GonoVac compared with 4CMenB in the current study both corroborates our mouse gonococcal challenge study results and highlights potential key immune mechanisms which may serve as the basis of protection.

Despite aluminum adjuvants typically promoting Th2 responses, GonoVac formulated with Al(OH)_3_ induced significant levels of IFNγ- and IL-17A-producing splenocytes. Similar findings have been observed in OMV-based vaccines targeting *E. coli* and *Klebsiella pneumoniae* (50, 51). IFN-γ is a key effector cytokine in host defense against intracellular and facultatively intracellular bacteria, promoting macrophage activation and enhancing antimicrobial functions (52, 53). In contrast, although IL-17A promotes neutrophil recruitment, studies in the murine gonococcal infection model have shown that IL-17A-dominated responses contribute to immune regulation that suppresses protective Th1- and Th2-mediated adaptive immunity (54, 55).

Considering the ability of gonococci to survive and sometimes replicate within phagocytes (56, 57), IFN-γ mediated induction and activation of phagocytic cells could be important for clearance of gonococcal infection. The importance of cellular immunity in controlling gonococcal infection is supported by intranasal and intravaginal immunization with gonococcal OMV co-administered with microencapsulated IL-12. These elicit robust Th1-driven responses and confer protection against gonococcal infection in the vaginal mouse infection model (58). These observations suggest the potential of adjuvants other than Al(OH)_3_ in enhancing efficacy of gonococcal OMV-based vaccines. The immunogenic equivalence of GonoVac produced in shake-flasks compared with bioreactors supports process scalability, critical for commercial manufacturing.

Despite these encouraging results, there are several limitations. First, despite GonoVac inducing robust and functional anti-gonococcal immunity, the correlates of protection against gonorrhea are unknown, and ultimately protective efficacy needs to be assessed in humans. Second, there are currently no international standardized gonococcal immunoassays or reference sera for calibration, making comparison of findings between studies, for example, with SBA, difficult. Third, although ELISpot assays revealed cytokine-secreting cell responses following immunization with GonoVac, the method does not distinguish between specific cellular subsets. Thus, while increases in IFN-γ- and IL-17A-producing splenocytes indicate Th1 and Th17 involvement, these cytokines may also be produced by other immune cells (59). Future studies employing intracellular cytokine staining or single-cell RNA sequencing will be necessary to delineate the specific cellular contributors to the observed cytokine patterns.

In conclusion, GonoVac induces potent and multifaceted immune activation in mice and rabbits spanning systemic and mucosal IgG, antibody functionality, and cellular responses. Although 4CMenB has demonstrated moderate cross-protection against gonorrhea and is now recommended in the UK to protect specific populations, the vaccine is derived from *N. meningitidis* and its primary target is *N. meningitidis*. By contrast, GonoVac is specifically formulated to target gonococcal antigens, potentially offering more robust and targeted immunity. We hypothesize that the enhanced gonococcal antigenic specificity, which this study demonstrates translates to superior gonococcal-specific immunogenicity, will enhance cross-reactivity and cross-protection across diverse gonococcal strains, addressing potential limitations associated with using meningococcal-derived vaccines such as 4CMenB to protect against gonorrhea.

These findings support further development of GonoVac, assessment of cross-reactivity of immunity against a panel of representative global gonococcal isolates, and assessment in clinical trials. Evaluating the vaccine with and without Al(OH)_3_, in humans will provide key insights into whether differences in immune profile of the two formulations in animals translate to humans. Such studies are essential to determine the optimal formulation for clinical use of GonoVac, in the context of rising antimicrobial resistance and the urgent need for effective gonococcal vaccines.

## Materials and Methods

### Strains and growth

*N. gonorrhoeae* GC_0817560 was obtained from the WHO Collaborating Centre for Gonorrhoea and other Sexually Transmitted Infections, Örebro, Sweden. *lpxL1* was deleted from the GonoVac nOMV-producing strain, GC_0817560*lpxL1^-^*, to reduce reactogenicity as previously described (28). Frozen bacterial glycerol stock was used to inoculate gonococcal agar (GCA) supplemented with 2% Vitox (Oxoid) (Becton Dickinson, UK) and incubated overnight at 37 °C with 5% CO_2_. Growth in liquid was performed in gonococcal broth (15 g/L proteose peptone no. 3 (Oxoid), 4 g/L K_2_PO_4_ (Sigma-Aldrich), 1 g/L KH_2_PO_4_ (Sigma-Aldrich) and 5 g/L NaCl (Sigma-Aldrich); pH 7.2 - sterilized by autoclaving) supplemented with 1% Kellogg’s supplement I (400 g/L D-glucose (Sigma-Aldrich), 10 g/L L-glutamine (Sigma-Aldrich) and 0.002% w/v thiamine pyrophosphate (Sigma-Aldrich) - sterilized by 0.2 µM filtration)(60). Liquid cultures were incubated at 37 °C with 5% CO_2_ and 170 rpm shaking.

### Production of gonococcal nOMVs in shake-flasks

Strains were grown in two consecutive 50 mL cultures in 250 mL baffled-vented shake flasks, until OD_600_ 0.6, before a third culture of 240 ml was inoculated in a 2 L flask. This culture was grown overnight at 37 °C with 5% CO_2_, then centrifuged at 4,500 x g for 20 min and the supernatant filtered using a 0.22 μm membrane to remove bacteria. The supernatant was ultracentrifuged at 31,000 g for 3 h and the pelleted OMV resuspended in phosphate-buffered saline (PBS-Sigma-Aldrich) and filtered using a 0.2 μm filter. nOMV total protein content was assessed by Bicinchoninic acid assay kit (BCA Protein Assay Kit Pierce, Thermo Scientific).

### Production of gonococcal nOMVs in Bioreactor

Frozen bacterial glycerol stock was propagated first in 25 mL, then in150 mL liquid culture in baffled-vented shake flasks, at 37 °C with 5% CO_2_ until OD_600_ 0.7-1.0. The whole volume of the second culture was used to inoculate 1.35 L of media in a stirred tank reactor (STR). The fermentation was performed for 16 hours at 37°C, pH 7.2 controlled by addition of NaOH, air flow at 1.5 L/min and impeller speed 100-700 RPM controlled by dissolved oxygen set at 90% of the initial value. Following host cell nucleic acid digestion, the culture was harvested with initial clarification over two sets of depth filters (diatomaceous earth and cellulose/polyethersulfone; 10/0.8µm and 0.22µm respectively) at a flow rate equal to 380 LMH and a volumetric challenge of 18 L/m^2^. Tangential flow filtration, on polyether sulfone hollow fiber (0.5 mm lumen) operated at 5000-1s shear rate, and 4 psi feed pressure, was used to concentrate the product 15-fold (volumetric challenge 63 L/m2 of 300kDa membrane) and to diafilter it using seven diafiltration volumes of phosphate-buffered saline (PBS-Sigma-Aldrich). The product was filtered on 0.22 µm PVDF membrane prior to vialing and placing into -80°C for storage.

### Production of gonococcal lysates

*N. gonorrhoeae* GC_0817560 wildtype liquid cultures were grown as previously described and harvested by centrifugation at 4,700 g for 15 min, the pellet resuspended in PBS and cells homogenized using Lysing Matrix B tubes (MP Biomedicals FastPrep-24) twice at 4 m/s for 40 sec with 5 min incubation on ice in between. Debris was pelleted at 10,000 g for 10 min. The supernatant was 0.22 µm filter-sterilized.

### LOS Extraction

*N. gonorrhoeae* GC_0817560 was propagated on agar plates as previously described. Bacterial cells were harvested from agar by scraping, resuspended in PBS and centrifuged at 10,000 rpm for 15 min at 10°C. The cell pellet was treated with a shearing buffer containing 2% v/v EDTA, 0.5% w/v sodium azide in PBS for 15 minutes at 60°C, and passed through 20-gauge needles to lyse cells and shear genomic DNA. The sheared suspension was treated with 10 µg/mL DNase and RNase at 37°C for 3 hours, and with 20 µg/mL of Proteinase K at 60°C for 3 hours. LOS was extracted by adding preheated phenol in 1:1 ratio and incubating at 60°C for 15 minutes with constant mixing, followed by centrifugation at 10,000 rpm for 15 minutes at 10°C. The upper LOS-containing aqueous layer was carefully decanted, and the extraction process was repeated by adding ultrapure water in a 1:1 ratio to the remaining bottom layer. The two LOS-containing aqueous layers were pooled and dialyzed with ultrapure water for 2 days with frequent water changes to remove phenol residues.

Further purification was performed by precipitation with 50 mM sodium acetate in 80% ethanol solution at -20°C overnight, followed by centrifugation at 5000 rpm for 15 minutes at 10°C. Following overnight drying at 37°C, the precipitate was resuspended in ultrapure water, ultracentrifuged at 30,000 rpm for 3 hours at 4°C and the pellet resuspended in ultrapure water. A final DNAse, RNase, and Proteinase K treatment was repeated as described previously. Purity of extracted LOS was assessed by SDS-PAGE with Coomassie blue staining and quantity assessed by SDS-PAGE with serially diluted *E. coli* LPS standard of known concentration following by silver staining and densitometric bands analysis.

### Animal immunogenicity studies

BALB/c mice (female, 6-8 weeks old, 6 mice per group) were housed in pathogen-free conditions and immunized as described in the result section. Intermediate blood samples were collected through tail bleeds and, terminal samples by cardiac puncture. Blood was allowed to clot then spun twice at 13,000 g for 10 min to separate the serum, which was stored at -40°C. Vaginal lavages were collected following three immunizations by flushing the vaginal canal with 30 µL PBS three times (total 90 µL) and mucosal anti-gonococcal IgG levels quantified by ELISA.

Spleens were dissected post-mortem and processed into single-cell suspensions using gentleMACS Dissociators and C tubes containing autoMACS Running Buffer (Miltenyi Biotec) as previously described (61). Cells were centrifuged (300 g, 10 min), treated with Red Blood Cell (RBC) lysis buffer (Qiagen) for 5 min, and the reaction quenched with autoMACS Rinsing Buffer. After further washing and filtration through 40 µm strainers (Greiner Bio-One), cells were resuspended in Recovery Cell Culture Freezing Medium (1–8 × 10⁷ cells/mL; Gibco), aliquoted into cryovials, and frozen at –80 °C using a CoolCell container (Corning) before transfer to liquid nitrogen for long-term storage. Rabbit studies were performed by Pharmaron under GLP-compliant conditions.

### Ethics statement

Mouse experiments were performed under UK Home Office Animals Act Project License PP7685090 and received approval from the University of Oxford Animal Care and Ethical Review Committee.

### ELISA to evaluate serum IgG, IgG subclasses, IgA and IgM and vaginal mucosal IgG and IgA

Indirect ELISA were developed to quantify gonococcal-specific immunoglobulins (IgG, IgG subclasses, IgA) and vaginal IgG. Nunc MaxiSorp 96-well plates were coated with 50 μL/well of GC_0817560 lysate (6 μg/mL in PBS) and incubated overnight at 4°C. After five washes with 0.05% PBS-Tween-20, wells were blocked with 200 μL 1% PBS-BSA for 2 h at 37 °C. Following additional washes, 50 μL of serially diluted standard curve reference sera, test samples (sera or vaginal lavages), and positive control sera (diluted in PBS-BSA) were added. Standard curves were generated using two-fold series in duplicate, with two blank wells as zero points. Plates were incubated with serum for 2 h at room temperature or overnight at 4 °C, washed, and incubated at room temperature with 50 μL/well HRP-conjugated secondary antibodies for 2 h.

The secondary antibodies were: 1:10,000 goat anti-mouse IgG-HRP (Jackson ImmunoResearch); 1:8,000 goat anti-mouse IgG1-, IgG2a-, IgG2b-, IgG3-, or IgM-HRP (SouthernBiotech); and 1:3,000 goat anti-mouse IgA-HRP (SouthernBiotech). After washing, 100 μL of TMB substrate was added until the highest standard reached OD_630_ 0.9, or, for IgA ELISA, for a fixed time of 20 min. The reaction was stopped by addition of 50 μL 2 M H₂SO₄. Standard curves were established using pooled immune sera from GonoVac-immunized mice, and positive controls with high, medium, and low antibody titers from *N. gonorrhoeae* FA1090 OMV-immunized mouse sera. Antibody concentrations were interpolated using a 4-parameter logistic model in Gen5 software, expressed in Arbitrary ELISA Units (AEU), and adjusted for sample dilution. Serum IgG titers were normalized by dividing by 100, the minimum dilution factor.

Mucosal IgA was assessed using Nunc MaxiSorp 384-well, round-bottom plates. Volumes were adjusted to 20 μL/well for coating antigen, primary, and secondary antibodies. Plates were blocked with 80 μL/well of 1% PBS-BSA, and the above procedure was followed.

### ELISA to evaluate rabbit serum anti-gonococcal IgG

Anti-gonococcal IgG titers in rabbit serum were determined by end-point ELISA. Briefly, 96-well Nunc Maxisorp plates were coated with GC_0817560 lysate using the method previously described. Rabbit sera were serially diluted two-fold in PBS-1% BSA starting at 1:6,400. Diluted sera were added to coated plates in duplicate and incubated for 2 h at room temperature. After washing, bound IgG was detected using goat anti-rabbit IgG conjugated to horseradish peroxidase (Sigma A0545; 1:10,000 in PBS-1% BSA). Plates were developed for 20 min with TMB substrate, and the reaction stopped with 2 M H₂SO₄.

Absorbance was read at 450–630 nm using a Biotek ELx808 plate reader. End-point titers were defined as the reciprocal of the last serum dilution giving an optical density (OD₄₅₀–₆_3_₀) at least three standard deviations above the mean of the negative control (naïve serum).

### Anti-LOS serum IgG ELISA

Anti-LOS serum IgG was assessed by in-house ELISA. 96-well Maxisorp plates were coated with 6 μg/mL of gonococcal LOS in PBS overnight at 4 °C. After washing, 50 μL of serially diluted standard reference sera, test samples (in duplicate), and positive control sera (in triplicate) were added. Following incubation, plates were washed, and 50 μL/well of a 1:5,000 dilution of goat anti-mouse IgG HRP-conjugated secondary antibody was added and incubated for 2 h. The subsequent steps followed the protocol described above.

### Serum IgG avidity

Avidity of serum anti-gonococcal IgG was assessed by gonococcal ELISA (above) using urea as chaotropic agent. Test sera were added at a single dilution in duplicate wells twice, one duplicate on each half of the plate. After 2 h incubation at room temperature or overnight incubation at 4°C, plates were washed 3 times. Serum samples on one half of the plate were incubated with 50 µL/well of 6 M urea (Sigma-Aldrich) in PBS for 30 min to disrupt hydrogen bonds and van der Waals forces. Standard curve serial dilutions, positive controls and the other duplicate wells containing serum test samples were incubated with PBS alone. ELISA plates were then washed 3 times, and the assay completed as described above. Avidity index of each sample was expressed as the percentage ratio between the titer of the sample treated with a chaotropic agent (urea) and the non-treated sample IgG titer.

### Serum bactericidal assay

*N. gonorrhoeae* GV_0817560 was grown in GCB until mid-log phase, then diluted in bactericidal buffer (Hanks’ Balanced Salt Solution (HBSS, Sigma-Aldrich) containing 0.5% BSA Sigma-Aldrich) to 6 x 10^4^ colony-forming units (CFU)/mL. Mouse test sera were heat-inactivated at 56 °C for 30 minutes. In a 96-well plate, 10 µL of the bacterial suspension and 10 µL of human complement (final assay concentration of 12.5%; IgG/IgM/IgA-depleted serum from Pel-Freez Biologicals) were added to wells containing ten two-fold serial dilutions of test serum samples in bactericidal buffer. The mixtures were incubated for 45 min at 37 °C. Post-incubation, 10 µL/well were plated on GCA to determine bacterial viability. Percentage survival was calculated relative to control wells containing bacteria and complement without sera, which represented 100% survival. The SBA titer was the highest serum dilution resulting in 50% bacterial killing compared to the CFU in the active complement no-sera controls.

### Cellular immune responses

Cellular immune responses were assessed by quantifying cytokine-secreting cells using a mouse FluoroSpot kit (Mabtech, FSP-4144-10). Frozen splenocytes were rapidly thawed in a 37°C water bath, then transferred to 10 mL R10 RPMI (Gibco) with 10% heat-inactivated fetal Calf Serum (FCS) (Sigma-Aldrich), 1% L-glutamine (Sigma-Aldrich), 1% Penicillin-Streptomycin (pen/strep; Sigma-Aldrich)) pre-warmed at 37°C and were washed with R10 by repeated centrifugation at 300 x g for 10 min. Pelleted cells were resuspended in 1 mL media and rested for 1 h at 37°C and 5% CO_2_ before pelleting by centrifugation for 5 min at 300 g, resuspended in 10 mL R10 and counted using a 1:1 dilution with trypan blue using the Countess II FL Automated Cell Counter (Invitrogen).

Cells were then diluted in 1 mL R10 to 2.5 x 10^6^ cells/mL and added to Fluorospot plates pre-coated with anti-IFN-γ, anti-IL4 and anti-IL17A monoclonal antibodies at 15 μg/ml (anti-IFNγ and anti-IL4) and 10 μg/ml (anti-IL-17A), respectively (Mabtech). 75 μg/mL (final concentration) of gonococcal GC_0817560 lysates and 0.2 µg/mL final concentration CD28 as co-stimulus were used in triplicate wells to assess gonococcal antigen-specific splenocytes (100 μL/well). Positive control stimulus (1 well per sample) contained 25 ng/mL phorbol 12-myristate 13-acetate (PMA, Cayman Chemical) and 500 ng/mL ionomycin (IO, Sigma-Aldrich). 50 μL PBS and 50 μL media were used in negative control wells, unstimulated, in duplicate. Cells were stimulated for 24 h, and spots counted the following day using AID ELISpot software 8.0. Results were the average of triplicate wells with subtraction of unstimulated controls.

### Dose-response curve

The dose-response analysis was performed using GraphPad Prism 10 using log-transformed antibody concentrations which were used to fit 4-parameter and 5-PL models. The model with the best fit was selected based on the R-squared adjusted value (adjuster R^2^), indicating the proportion of variance explained by the model and the Root Mean Square Error (RMSE), both determined by the GraphPad software for each data set. The adjR^2^ is a version of the coefficient of determination adjusted to take into account number of predictors in the model with high adjusted R^2^ suggesting adequate fitting (33). RMSE quantifies the average deviation between predicted and observed values, with lower values reflecting improved accuracy (34, 35).

### Statistical analyses

Statistical analyses were conducted using GraphPad Prism (version 10) and R statistical software (version 4.5.1) (62). All numerical datasets were assessed for normality on both original and log-transformed scales to guide the choice between parametric and non-parametric methods. Depending on distributional assumptions, group comparisons were performed using one-way analysis of variance (ANOVA) or the Kruskal–Wallis test, followed by appropriate post hoc multiple-comparison procedures (Tukey’s or Dunn’s tests, respectively). Two-way ANOVA with Šidák adjustment for multiple comparisons was applied when evaluating the effects of two independent factors. Prior to two-way ANOVA, ELISA and SBA measurements were log-transformed (log10 and log2, respectively) to reduce heteroscedasticity or minimize data variability and improve model assumptions. Bivariate associations between underlying numerical variables were evaluated using the non-parametric Spearman’s rank correlation coefficient (*r*) implemented via the PerformanceAnalytics R package (63).

## Acknowledgements

This work was funded by CARB-X (grant number 06CARB-X0948). The funders had no role in study design, data collection and interpretation, or the decision to submit the work for publication.

We thank Professor Magnus Unemo, WHO Collaborating Centre for Gonorrhoea and other Sexually Transmitted Infections, Örebro, Sweden, for provision of *N. gonorrhoeae* GC_0817560.

